# Role of ischemic preconditioning in cardioprotective mechanisms of mCRP deposited myocardium in a rat model

**DOI:** 10.1101/2020.05.13.093989

**Authors:** Eun Na Kim, Jae-Sung Choi, Chong Jai Kim, So Ra Kim, Ki-Bong Kim, Se Jin Oh

## Abstract

The deposition of monomeric C-reactive protein (mCRP) in the myocardium aggravates ischemia-reperfusion injury (IRI) and myocardial infarction. Ischemic preconditioning (IPC) is known to protect the myocardium against IRI. We evaluated the effects of IPC on mCRP-deposited myocardium due to IRI in a rat model. Myocardial IRI was produced by ligation of the coronary artery. Direct IPC was applied before IRI using multiple short direct occlusions of the coronary artery. CRP was infused intravenously after IRI. The study groups included the following: sham (n=3), IRI only (n=5), IRI+CRP (n=9), and IPC+IRI+CRP (n=6) groups. The infarct area and area at risk were assessed using Evans blue and 2,3,5-triphenyltetrazolium chloride (2,3,5-TTC) staining. Additionally, mCRP immunostaining and interleukin (IL)-6 mRNA reverse transcriptase-polymerase chain reaction (RT-PCR) were performed. In the IRI+CRP group, the infarcted area, mCRP deposition, and IL-6 mRNA expression were higher than those in the IRI only group. However, in the IPC+IRI+CRP group, the infarction (20% vs. 34% p=0.085) and mCRP myocardial deposition (21% vs. 44%, p=0.026) were lower and IL-6 mRNA expression was higher than those in the IRI+CRP group (fold change, 407 vs. 326, p=0.808), although this was not statistically significant. IPC has cardioprotective effects against myocardial damage caused by mCRP deposition. This protective effect is related to the increase in IL-6 mRNA expression.

## Introduction

Ischemic heart disease is one of the most important causes of global death, causing over 7.2 million deaths in the world annually[1]. In order to reduce the myocardial infarct size and prevent poor clinical outcomes, coronary artery reperfusion of the ischemic myocardium is performed using thrombolytic therapy, percutaneous coronary intervention, or coronary arterial bypass grafting. However, when the ischemic myocardium is reperfused, it can result in paradoxical harmful effects that damage the myocardium—this is called ischemic reperfusion injury (IRI)[2]. Approximately 50% of the final size of myocardial infarction is due to IRI[1]. Subsequently, 10% of deaths and 25% of cardiac failure following acute myocardial infarction is due to IRI even after treatment with reperfusion of the ischemic heart[3]. Therefore, minimizing IRI is the most important strategy to salvage the myocardium after an ischemic event.

In 1986, Murry et al. proposed the concept of the cardioprotective role of ischemic preconditioning (IPC); multiple, brief, non-lethal ischemic episodes followed by short reperfusion before the main prolonged ischemic injury reduces the infarct size due to the development of resistance to IRI and inhibition of lethal reperfusion injury[4]. For decades, many researchers have studied the efficacy and mechanisms of IPC, and the myocardial protective effects of IPC have been demonstrated in animal studies[5, 6], *in vivo* human heart studies[7–9], and *in vitro* studies[10]. Ischemic preconditioning not only reduces the infarct size by increasing the resistance of isolated myocytes to hypoxic injury[11], but also reduces anginal pain, ST-segment elevation and lactate production[12], and reduces post-ischemic arrhythmia[13, 14]. IPC also slows metabolism[15] and helps in the recovery of cardiac function after an ischemic event[16].

There is emerging experimental evidence that the deposition of monomeric C-reactive protein (mCRP) exacerbates the damage to the heart due to IRI[17, 18]. CRP is an acute phase reactant protein mainly produced in the liver during systemic infections and inflammation[19]. The serum level of CRP is not only an important prognostic and predictive marker for various cardiovascular diseases[20] including the clinical outcomes, death, and heart failure following myocardial infarction[21] but CRP itself also acts as a cause of direct damage to the cardiovascular tissue[22, 23].

CRP in serum exists in a pentameric form (pCRP), and when it encounters a damaged cell membrane it undergoes structural changes from pCRP to mCRP[24]. Subsequently, mCRP is deposited in the damaged tissue, thereby, activating the reactive oxygen species[25] and complement system[26]. This aggravates the inflammatory process and exacerbates myocardial damage[18]. Using a rat IRI model, we previously confirmed that if the serum CRP level is high during the myocardial ischemic-reperfusion insult, serum CRP is deposited in the myocardium as mCRP and the size of the myocardial infarction is increased[17]. Additionally, our previous work demonstrated that the microRNA profile of the myocardial area at risk changed drastically when CRP was high during the ischemic-reperfusion injury[27].

However, the role of IPC in the myocardium damaged by IRI with mCRP deposition has not been investigated. Therefore, in this study, we investigated whether IPC is protective against mCRP-induced myocardial damage in IRI settings using a rat acute myocardial IRI model.

## Materials and methods

### Animals

We set up a myocardial IRI model using female Sprague–Dawley rats weighing between 220 and 270 g with a gestational age of 10–14 weeks. The animals were treated according to the *Guide for the Care and Use of Laboratory Animals* (National Academy of Sciences, Washington, DC, USA). The protocols for animal use were approved by the Institutional Animal Care and Use Committee (IACUC) at the SMG-SNU Boramae Medical Center Biomedical Research Institute (approval number: 2016-0027). Anesthesia was administered with inhalation of isoflurane (4%) for induction, followed by intraperitoneal administration of Tiletamine HCl and Zolazepam HCL (Zoletil 50^®^, 0.12 mL, Virbac, Carros, France) and xylazine (Rompun^®^, 2%, 0.02 mL, Bayer Healthcare, Loos, France) for maintenance.

The rats were intubated with 16-gauge intravenous catheters (REF 382457; BD Medical, Sandy, Utah, USA) and connected to ventilators (683 rodent ventilator; Harvard Apparatus, Holliston, Massachusetts, USA). Positive pressure ventilation at room air with a tidal volume of 2.5 ml (10 mL/kg, 60 breath/min) was used to prevent atelectasis during the procedure. We approached the heart through left thoracotomy via the fourth intercostal space. The pericardium was opened to expose the left coronary artery. Myocardial ischemic injury was produced by ligating the left anterior descending coronary artery (LAD) approximately 2 mm distal to the origin of LAD using 6-0 nylon double sutures, buttressed with a small piece of plastic tube (Fig 1). After 45 minutes of ischemia, we loosened the sutures to allow for reperfusion for 45 minutes. The pericardium was left open to expose LAD during the procedure.

**Fig 1.**
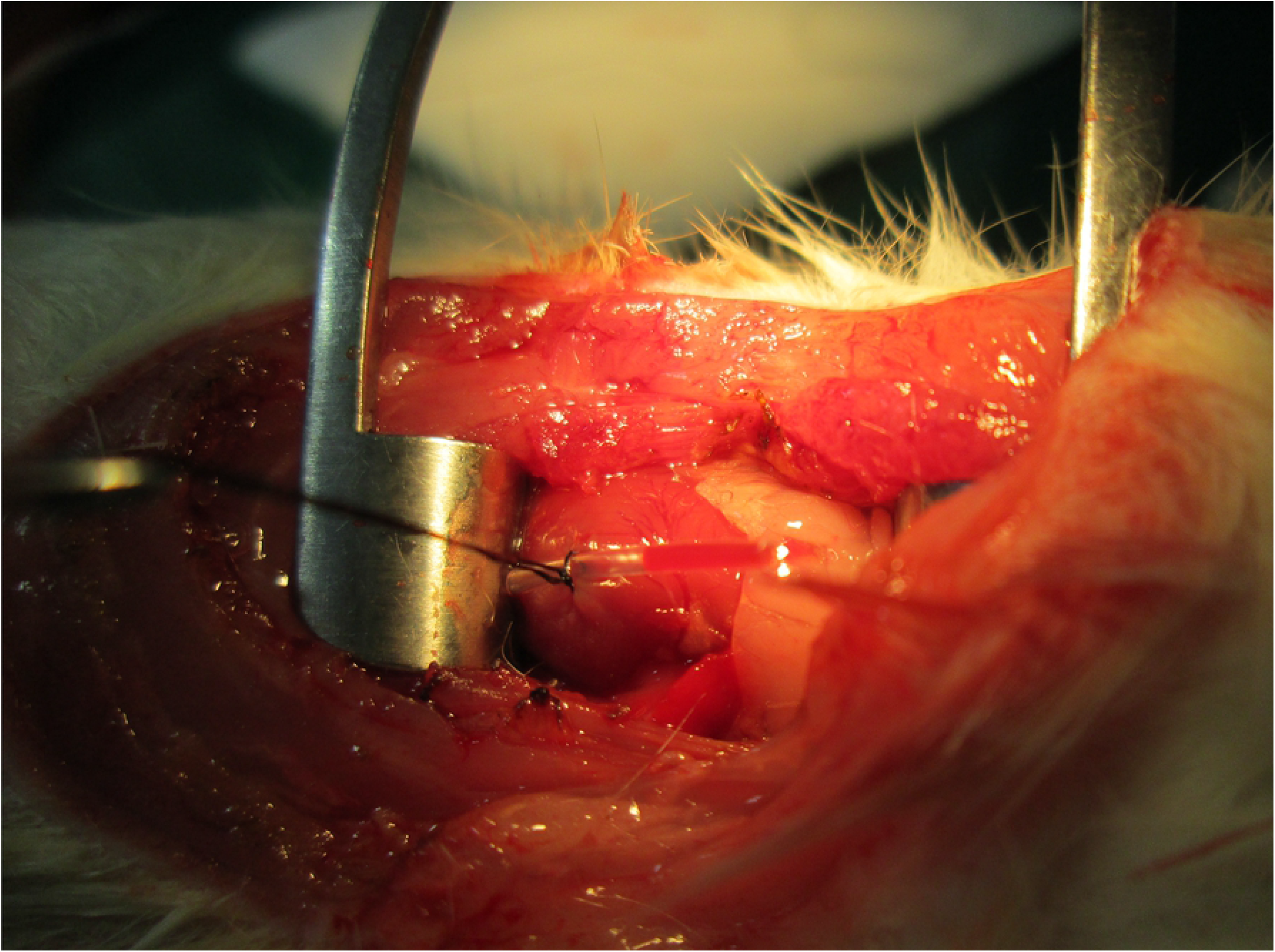
Ligation of the left anterior descending coronary artery, buttressed with a plastic tube.

### Experimental protocols

The experimental protocols are illustrated in Fig 2. In the sham group (n=3), thoracotomy and pericardiotomy were performed, and the pericardium was opened and maintained for 90 minutes without any manipulation. Subsequently, the rats were euthanized and autopsy was performed (Fig 2A). In the group with IRI only (IRI only group, n=5), the myocardium was excised quickly after 45 minutes of LAD ligation and 45 minutes of reperfusion (Fig 2B). In the group treated with CRP following IRI (IRI+CRP group, n=9), high-purity (>99%) human CRP obtained from human plasma (C4063; Sigma-Aldrich, Saint Louis, Missouri) was infused via the femoral vein after 45 minutes of ischemia with LAD ligation, just before reperfusion (Fig 2C). In the group treated with ischemic preconditioning (IPC) followed by IRI and CRP injection (IPC+IRI+CRP group, n=6), IPC was applied before LAD ligation; IPC included three occlusions for 3 minutes each followed by a 5-minute period of reperfusion after each occlusion (Fig 2D).

**Fig 2.**
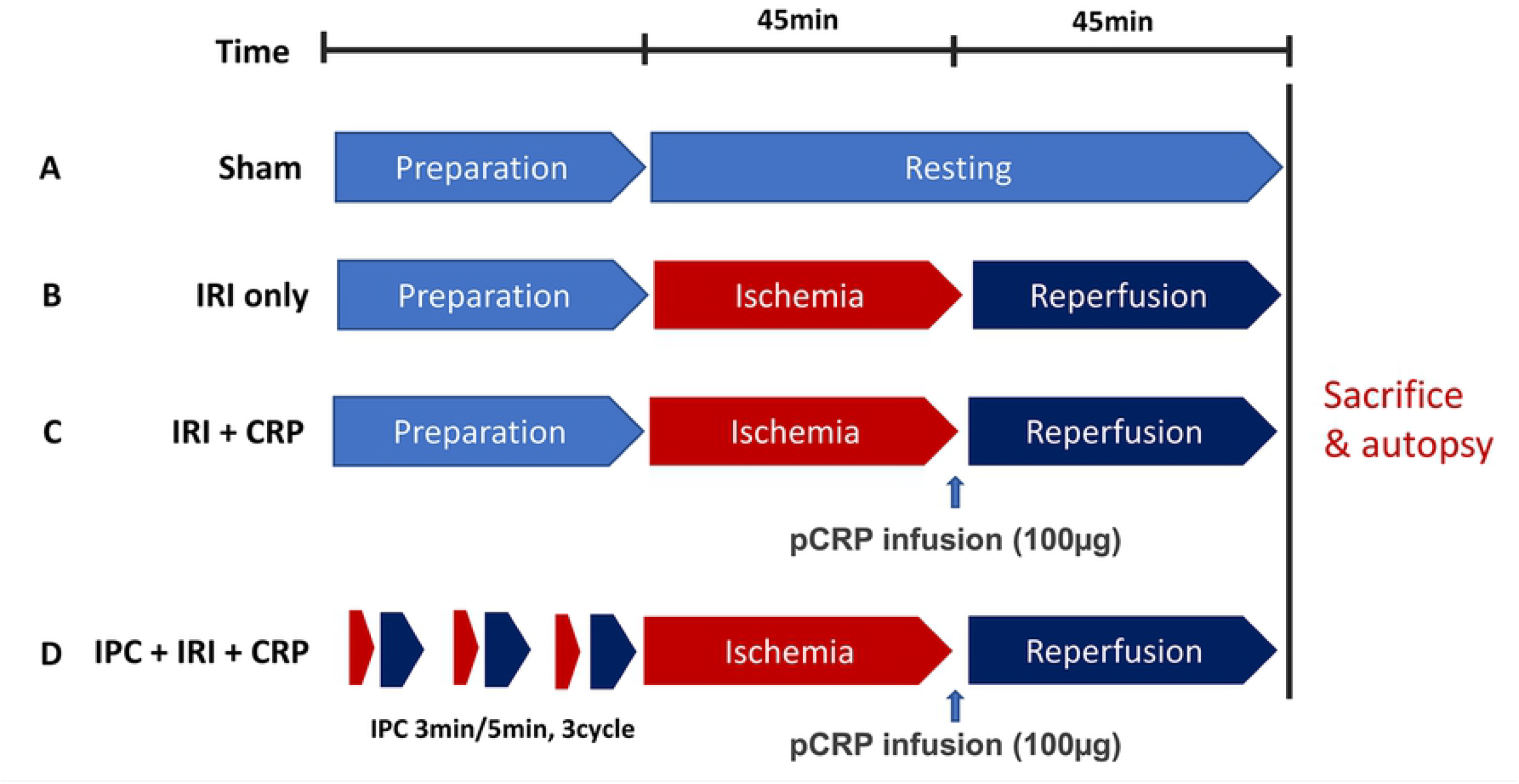
The experimental protocols.

### Evans blue and TTC staining to determine infarct area and AAR

To evaluate the non-ischemic area, ischemic but not infarcted viable area (area at risk, AAR), and infarcted area, we performed staining with Evans blue and 2,3,5-triphenyltetrazolium chloride (TTC, Sigma Chemical, Saint Louis, MO). After euthanasia and before the removal of the heart, 1 mL diluted heparin solution (2500 IU heparin/mL) was infused via the coronary ostia after clamping the ascending aorta. We re-sutured the LAD artery using 6-0 nylon, and injected 1% Evans blue solution to stain the perfused non-ischemic myocardium. Both AAR and infarcted area (whole ischemic area) do not stain with Evans blue solution[28]. After Evans blue perfusion, the heart was cut into four transverse sections at regular intervals from the apex to the base[29]. One of the middle sections was used to measure the ischemia and infarct size. This midportion was sliced again in 4-mm slices and one slice was incubated with TTC dissolved in 100 mmol/L of phosphate buffer for 15 minutes. With TTC staining, the viable area of the myocardium—the non-ischemic area—and the viable area at risk are stained deep red and the infarcted zone remains unstained and is, therefore, white. Therefore, double staining with Evans blue and TTC stains the infracted area white, AAR deep red, and non-ischemic area blue[30].

### Histopathologic analysis and Immunohistochemistry

The middle section of the excised heart was fixed in 10% buffered formalin and embedded in paraffin. The 4-μm tissue sections were stained using hematoxylin and eosin (H&E), and immunostained with human monoclonal anti-CRP antibody (C1688; Sigma-Aldrich, Saint Louis, MO, USA; 1:400 dilution) which specifically detect the 24-kD monomeric CRP epitope[31, 32]. OptiView DAB immunohistochemical detection kit (Roche Diagnostics, Mannheim, Germany) and a Benchmark XT autoimmunostainer (Ventana Medical Systems, Tucson, AZ, USA) were used for immunostaining. Thorough histopathologic examination with microscopy was performed by one pathologist (E.N.K).

### Image analysis

Images of the heart specimens stained with Evans blue and TTC were captured with a digital camera DP26 (Olympus, Tokyo, Japan). Digital images of mCRP immunostaining were acquired using the Vectra automated imaging system (PerkinElmer, Waltham, MA, USA). The areas of the infarcted myocardium (white zone) and AAR (red zone) were automatically calculated with inForm (PerkinElmer) imaging analysis software. The size of the infarct was expressed as a percentage of the infarct area and the whole ischemic area (infarct / [infarct + AAR] × 100). mCRP immunolabeling was examined to determine the tissue distribution within each section. mCRP immunopositivity were expressed as the percentage of the mCRP-immunopositive area and the whole ischemic area (mCRP-immunostained area / [infarct + AAR] × 100) as previously reported[17].

### Analysis of IL-6 mRNA expression assay in rat myocardium

RNA was prepared using a miRNeasy Mini Kit (Qiagen, Hilden, Germany) according to the manufacturer’s instructions. The extracted RNA (1 μg) was reverse-transcribed using Reverse Transcription System (Promega, Madison, Wisconsin, USA), and cDNAs were amplified using GeneAmp PCR System 9700 (Applied Biosystems, Foster City, CA, USA). Quantitative reverse transcriptase-polymerase chain reaction (qRT-PCR) analysis of interleukin (IL)-6 was performed using TaqMan Gene Expression Assays (Rn01410330_m1; Applied Biosystems, Foster City, CA, USA) and 7900HT Fast Real-Time PCR System (Applied Biosystems). The rat ACTB (Rn00667869_m1; Applied Biosystems, Foster City, CA) endogenous control was used for normalization.

### Statistical analysis

Data was expressed as mean and standard deviation and plotted as mean with standard error of mean. To compare two groups, two-tailed *t*-test was used. A *P*-value < 0.05 was considered statistically significant. Data analyses were performed using GraphPad Prism v5.0 (GraphPad Software, San Diego, CA, USA).

## Results

### Areas of infarcted and ischemic myocardium

After Evans blue and TTC staining, the non-ischemic area was stained blue, AAR was stained red, and infarct area was stained white, as depicted in Fig 3. As with our previous study[17], the areas of myocardium damaged by IRI demonstrated cellular changes along with contraction bands with intensely eosinophilic intracellular stripes on H&E staining, thus confirming that IRI was performed well in the experiment[33].

**Fig 3.**
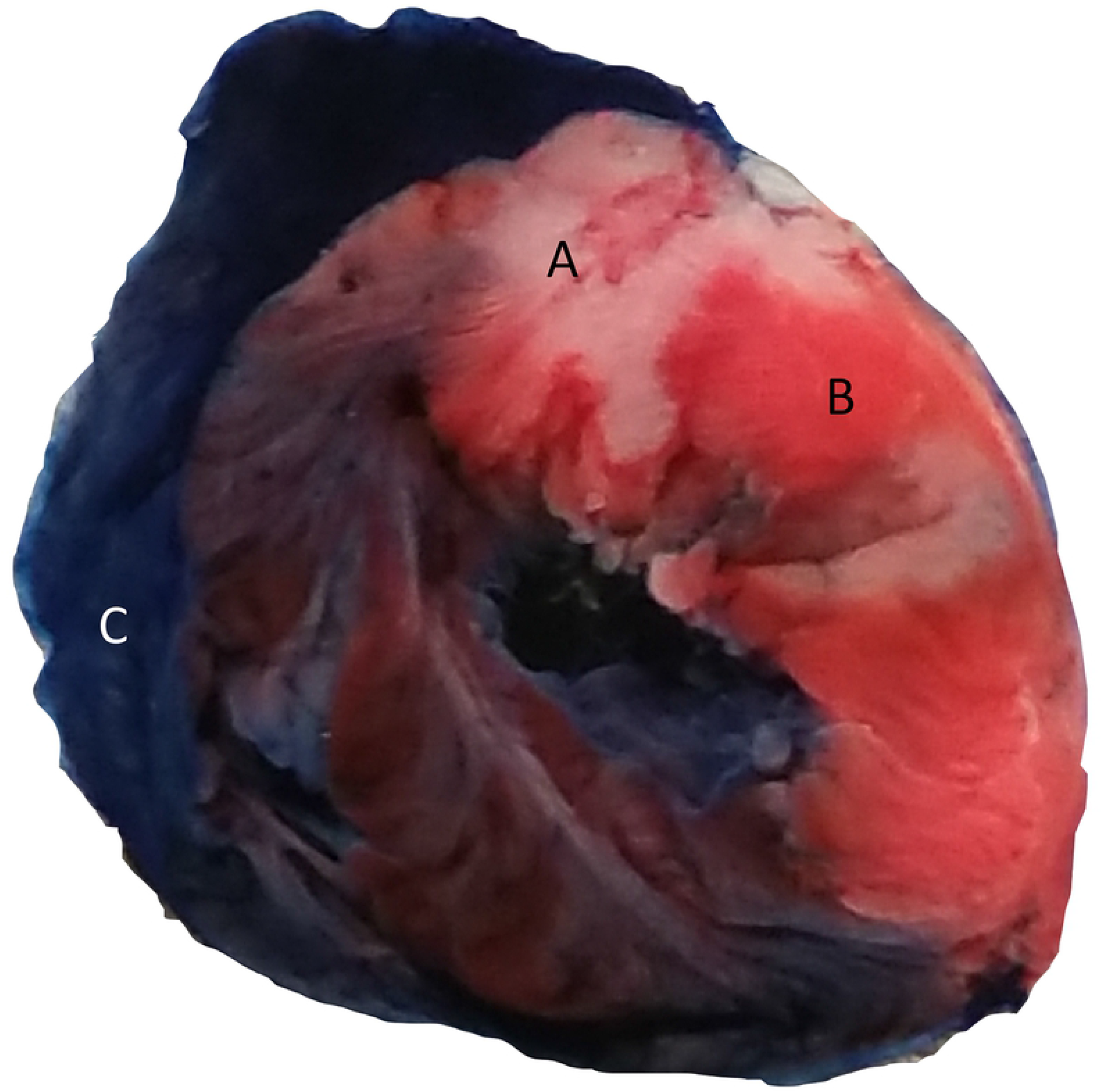
Evans blue and 2,3,5-triphenyltetrazolium chloride (TTC) staining. A. White area: infarcted area B. Red area: area at risk (ischemic but not infarcted) C. Blue area: non-ischemic area

The percentage of infarct area and whole ischemic area (infarct / [infarct + AAR] × 100) tended to be higher in the IRI+CRP group than that in the IRI only group (34±15% vs. 23±7%, *p* 0.116). After ischemic preconditioning, the size of the infarcted area decreased more in the IPC+IRI+CRP group than that in the IRI+CRP group, although statistical significance was not reached (21±7% vs. 34±15%, *p*=0.085, Fig 4A, 5).

**Fig 4.**
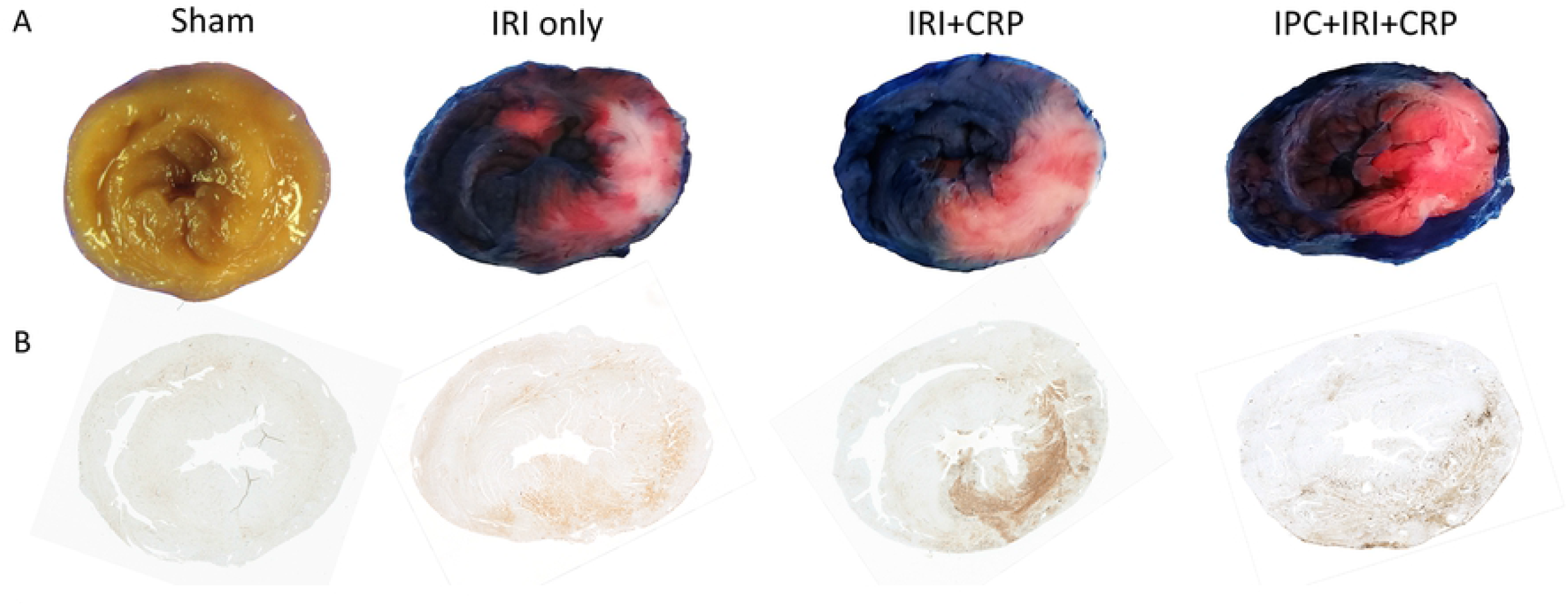
(A) The sham group without staining. TTC and Evans blue staining in the IRI only, IRI+CRP, and IPC+IRI+CRP groups. (B) mCRP immunohistochemistry in the sham, IRI only, IRI+CRP, and IPC+IRI+CRP groups. mCRP, monomeric C-reactive protein; IRI, ischemia reperfusion injury; TTC, 2,3,5-triphenyltetrazolium chloride.

**Fig 5.**
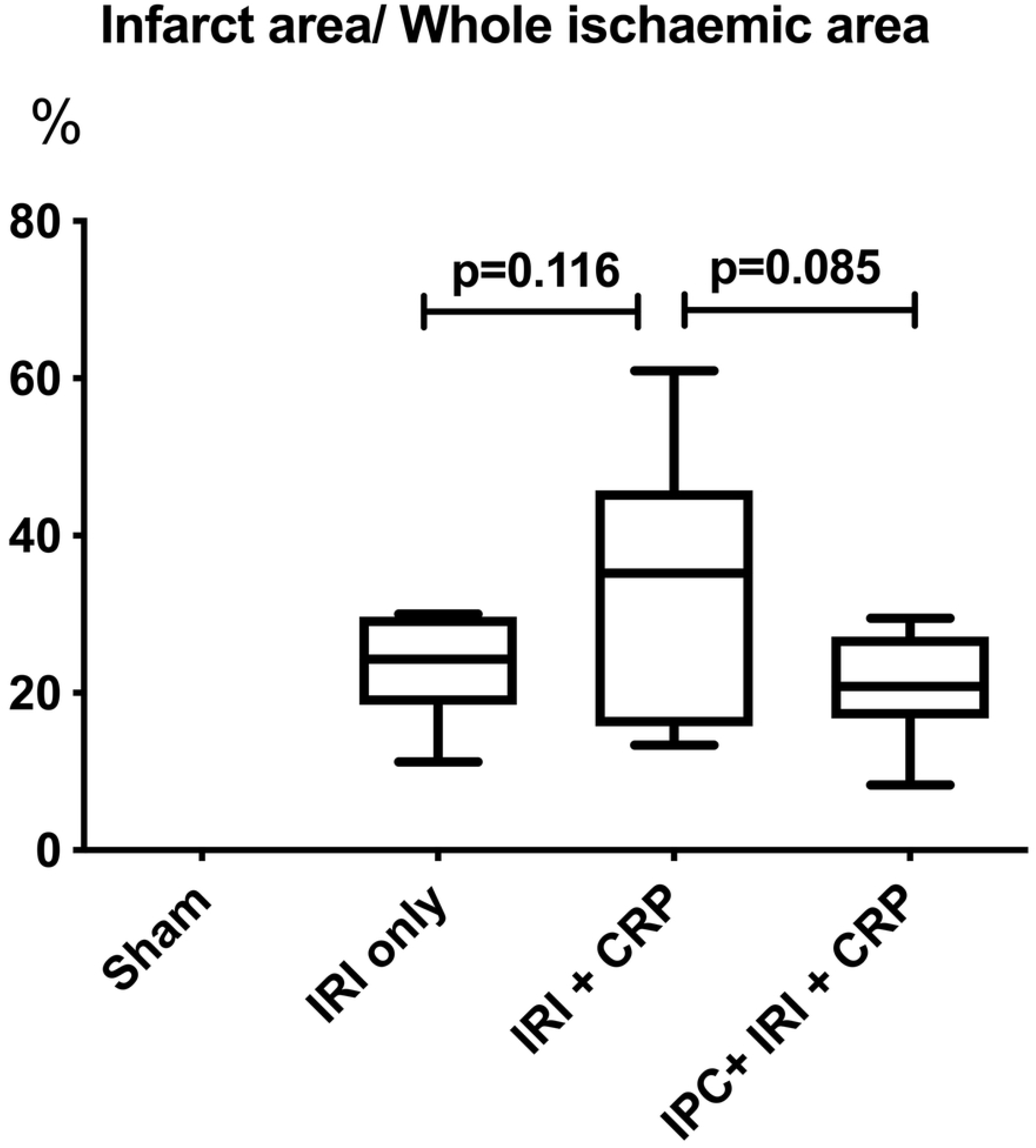
The ratio of infarct area and whole ischemic area.

### mCRP immunohistochemistry

While mCRP staining was faint and non-specific in the sham and IRI only groups, floating serum CRP was strongly and diffusely deposited on the damaged myocardium in the IRI+CRP group, not only in infarcted area, but also the viable AAR. This finding is consistent with our previous results[17](Fig 4B).

However, after applying direct IPC, the area of mCRP deposition in the ischemic myocardium (mCRP-immunostained area / [infarct + AAR] × 100)) was significantly lower than that in the IRI+CRP group (21 ±16% vs. 44±19%, p= 0.026, Fig 4B, 6).

**Fig 6.**
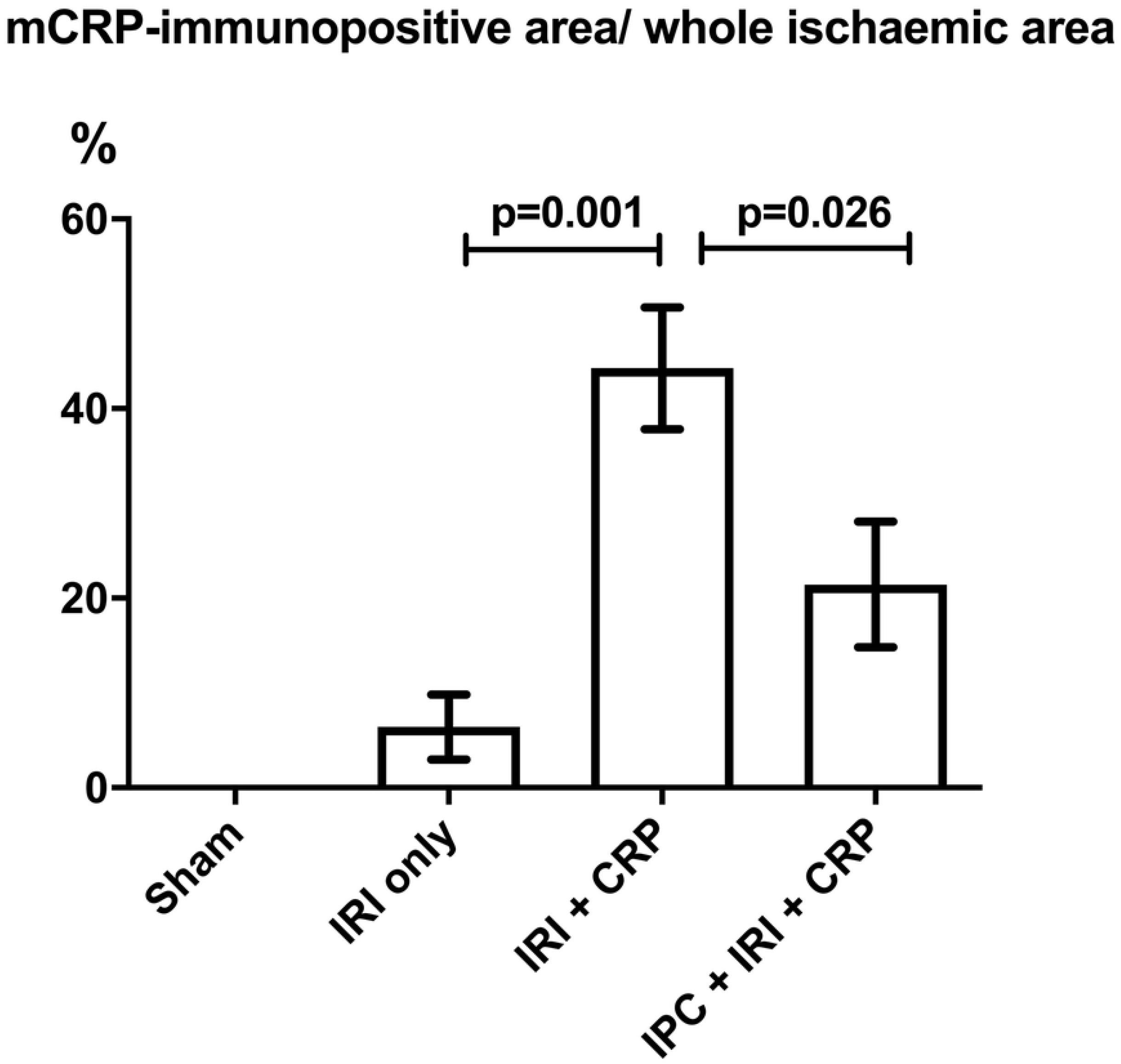
The ratio of mCRP-immunopositive area and whole ischemic area. mCRP, monomeric C-reactive protein.

### IL-6 mRNA expression increased after CRP injection and IPC

The IL-6 mRNA expression level was highest in the IPC+IRI+CRP group, followed by the IRI+CRP and IRI only groups (fold change, IPC+IRI+CRP group, 408±273, IRI+CRP group, 326±157, IRI only group, 198±113, Fig 7).

**Fig 7.**
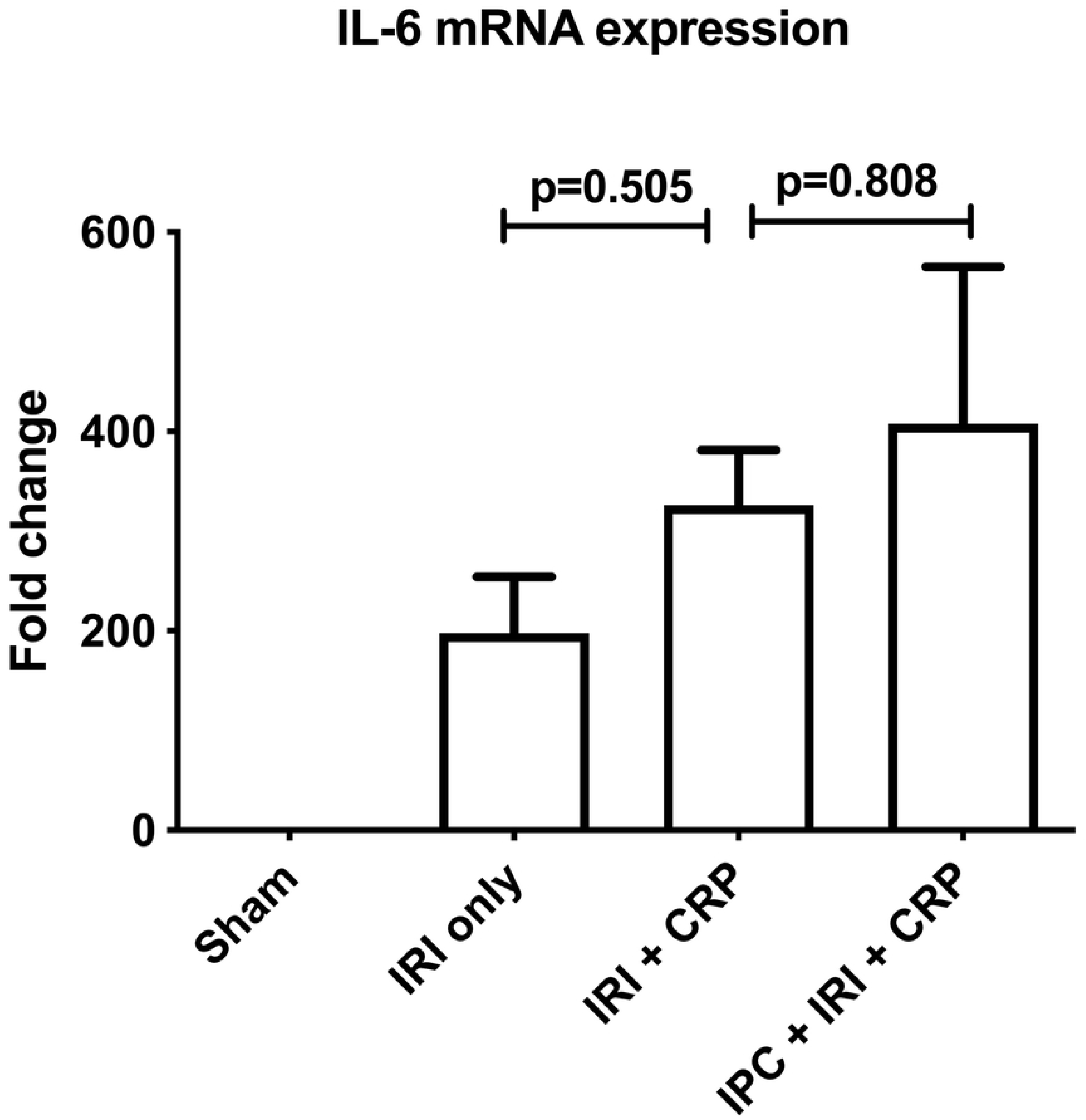
IL-6 mRNA expression. IL, interleukin; mRNA, messenger RNA.

## Discussion

To our knowledge, this is the first report of the cardioprotective effect of IPC on mCRP deposited ischemic myocardium. IPC-induced cardioprotective effects are known to be achieved by various triggers, receptors, and mediators[34]. Especially, adenosine, opioid receptors[35], bradykinin[36], and free radicals[37] play important roles in IPC. In a heart preconditioned with IPC, lactate accumulates slowly, which results in anti-infarct effects[15]. IPC reduces intracellular acidification and affects the sodium-proton exchanger, cytoskeleton changes, and down regulation of TNF alpha[34]. However, despite numerous studies over a long period of time, the comprehensive mechanism of IPC is still controversial and much remains unclear[4, 38, 39]. And, there have been no studies focused on the effects of IPC on mCRP-deposited ischemic and damaged myocardium; therefore, we investigated whether IPC affects the amount of mCRP deposition and myocardial infarction in IRI settings. In this study, we found that shortterm direct IPC prior to IRI and CRP-infusion reduces mCRP deposition in the myocardium and infarction size and increases IL-6 mRNA expression.

Previous works have documented that mCRP deposition aggravates IRI-induced myocardial infarction. Thiele et al. reported that in a rat model of IRI, mCRP was localized to the infarcted myocardium and aggravated inflammation via phospholipase A2-dependent dissociation of circulating pCRP to mCRP[18]. Pepys et al. reported that they used a 1,6-bis(phosphocholine)-hexane that can bind and inhibit human CRP to reduce the size of myocardial infarction[40]. Additionally, we previously reported that mCRP was deposited not only in the infarcted area but also in the area at risk along with mitochondrial damage and complement activation[17].

IL-6 is a pleiotropic pro-inflammatory cytokine and one of the main factors for the stimulation of acute phase proteins, such as CRP[41]. IL-6 is known to be released from cardiomyocytes under hypoxic conditions, such as myocytes in the border zone of myocardial infarction, and IL-6 derived from hypoxic myocytes may play an important role in the aggravation of myocardial dysfunction following IRI[42, 43]. Therefore, IL-6 was thought to result in hypertrophy and heart failure after ischemia by reducing the contractility of the myocardium[44]. In contrast, many studies have demonstrated that higher IL-6 levels during preconditioning play an organo-protective role[45–47]. Dawn et al. showed that ischemic preconditioning markedly upregulates IL-6 expression in the ischemic/reperfused myocardium. Furthermore, they demonstrated that IL-6 signaling plays an obligatory role in late preconditioning because IL-6 is required for the JAK-STAT signaling and upregulation of iNOS and COX-2, which are co-mediators of late preconditioning, and can, thus, aid in cardioprotection[45]. Similarly, Waldow et al. reported that if remote IPC was applied before IRI in a porcine lung, IL-6 increased more consistently than IRI alone, and the lung damage was alleviated[46]. In their *in vivo* study using ventricular cardiomyocytes from isolated rat hearts, Smart et al. demonstrated that IL-6-induced PI-3 kinase and NO-dependent protection of cardiomyocytes—associated with alterations in mitochondrial Ca^2+^ handling, inhibition of reperfusion-induced mitochondrial depolarization, swelling and loss of structural integrity, and suppression of cytosolic Ca^2+^ transients[47]. Additionally, in the present study, we confirmed that administering IPC before myocardial damage by IRI and mCRP deposition increases IL-6 mRNA expression and decreases the myocardial infarct size.

However, it is still controversial if increased IL-6 secretion plays an organo-protective role. In a porcine lung IRI model, Harkin et al. reported that IPC lowered IL-6 levels and resulted in a lung-protective effect[48]. The downstream pathways in which IL-6 plays a cardioprotective role are not well understood. Additionally, no detailed studies have been conducted to identify the specific signaling pathways when IL-6 is increased after mCRP deposition occurs. Further research is required to elucidate the mechanisms through which IL-6 acts on mCRP-deposited damaged tissue.

## Limitations

In the above experiment, some observations that did not demonstrate statistical significance were observed. However, when CRP was deposited, the infarction area increased with the same trend as that reported in our previous work[17]. Animal studies with a larger sample size are required to further validate these findings. There is a possibility of disparity between the findings in animals and real-world clinical situations. So far, numerous attempts to prevent IRI that worked in animal experiments have actually been ineffective in clinical studies[1]. We used an ischemia model that directly occludes healthy coronary artery in young rats, which is very invasive for practical clinical applications. It is also virtually impossible to predict when profound ischemia will occur and apply preconditioning before IRI. Therefore, it may be difficult to create a useful clinical application from our experimental results. Furthermore, to elaborate the animal model to reflect the actual clinical situations, animal models with older animals and comorbidities, such as diabetes, hyperlipidemia, atherosclerosis, and hypertension should be used[1].

There are many clinical situations in which serum CRP can be elevated in cardiac ischemia. Baseline CRP levels are moderately elevated in obese persons or those who smoke or have diabetes or hypertension. Additionally, CRP levels increase dramatically in patients with myocardial infarction[49]. If a patient with a cardiovascular event has elevated serum CRP level for any reason, the serum CRP will deteriorate myocardial function while getting deposited in the damaged myocardium. We, therefore, can infer that in these clinical situations, IPC will function to protect the mCRP deposited myocardium. Additionally, our experiments provide clues to understand the mechanisms of aggravation of ischemia due to mCRP and the protective mechanisms of ischemic preconditioning. If the mechanism by which IPC protects the myocardium that has undergone mCRP deposition is understood completely, it will provide a basis for developing a preconditioning mimetic agent.

## Conclusion

Ischemic preconditioning protects against myocardial damage caused by IRI and mCRP deposition. Additionally, this protective effect is believed to be associated with enhanced IL-6 mRNA expression.

## Acknowledgements

We thank Kyung Min Park, Gil-Je Lee and Yun Jae Kim for the image analysis, and Eun Jung Jeon for the molecular work. And we would like to thank Editage (www.editage.co.kr) for English language editing.

## Financial support

This work was funded by a clinical research grant-in-aid from the Seoul Metropolitan Government Seoul National University (SMG-SNU) Boramae Medical Center (03-2015-11). The funders had no role in study design, data collection and analysis, decision to publish, or preparation of the manuscript.

## Potential conflicts of interest

The authors have declared that no competing interests exist.

